# High consistency of trophic niches in soil microarthropod species (Oribatida, Acari) across soil depth and forest type

**DOI:** 10.1101/2021.07.24.453652

**Authors:** Jing-Zhong Lu, Peter Cordes, Mark Maraun, Stefan Scheu

## Abstract

Individuals of species may differ in resource use within and between populations. High intraspecific variation in resource use may hamper the co-existence of species in natural communities. To better understand the intraspecific variation in trophic niches of oribatid mites (Oribatida, Acari), we quantified stable isotope ratios of carbon (δ^13^C) and nitrogen (δ^15^N) of 40 Oribatida species that co-occurred in litter and soil of five forest types (European beech, Douglas fir, Norway spruce, two beech–conifer mixed forests) covering a range of environmental conditions. We found that although stable isotopes in litter and soil varied among forest types, δ^13^C and δ^15^N values of Oribatida and their trophic niches were remarkably stable between litter and soil, and also among forest types. We considered four trophic guilds of Oribatida representing the guild composition of the regional species pool; notably, trophic niches of Oribatida guilds also did not vary with soil depth. Furthermore, δ^13^C of Oribatida was more enriched (detrital shift) in European beech than in coniferous forests, but δ^15^N of Oribatida did not vary among forest types, indicating that basal resources of Oribatida are variable, but trophic positions are highly consistent across forest ecosystems. We conclude that trophic positions of Oribatida species and guilds are consistent across different forest types, and Oribatida species occupy virtually identical trophic niches irrespective of the soil depth they are colonizing. Overall, the results suggest that low intraspecific variability facilitates Oribatida niche differentiation and species coexistence.

## Introduction

In soil food webs, a large part of the isotopic space is occupied by soil microarthropods (Scheu and Falca, 2000; Pollierer et al., 2009; Maraun et al., 2011; Potapov et al., 2019). Based on the analysis of natural variations in bulk stable isotope values of carbon (^13^C) and nitrogen (^15^N), trophic niche differentiation has been uncovered in major groups of soil microarthropods including Oribatida, Collembola, and Mesostigmata (Schneider et al., 2004; Chahartaghi et al., 2005; Klarner et al., 2013). While trophic niche differentiation contributes to the co-existence of species, high intraspecific variation in stable isotope values points toward large niche overlap potentially hampering species co-existence (Schneider et al., 2004; Hart et al., 2016). For example, oribatid mites (Oribatida, Acari), typically occupy three to four trophic levels in temperate forests (Schneider et al., 2004; Maraun et al., 2011), but it remains poorly studied how trophic niches of Oribatida species vary with environmental conditions (Gan et al., 2014), such as soil depth and forest type.

Microarthropods dominate in the uppermost horizons of forest soils, especially the litter and the other organic horizons (Pande and Berthet, 1975; Mitchell, 1978; Arribas et al., 2021). Oribatida species were suggested to occupy similar trophic positions irrespective of soil depth (Scheu and Falca, 2000), but their variation in trophic niches with soil depth has not been rigorously tested. Stable isotope ratios of carbon and nitrogen are typically enriched in soil compared to litter (Potapov et al., 2019; Högberg et al., 2020), and if Oribatida species are generalist feeders (Maraun et al., 1998), their stable isotope values should increase with soil depth parallel to that in organic matter. Accordingly, if Oribatida species in soil incorporate older carbon than in litter, high stable isotope values should inflate their trophic position (Potapov et al., 2019). This may confound our understanding of their trophic ecology and their position in soil food web models. Variations in trophic niches of Oribatida species with soil depth may also be related to shifts in stable isotope values of the microorganisms digested. In organic matter of more advanced stages of decay deeper in soil, the proportion of bacteria and mycorrhizal fungi increases relative to saprotrophic fungi, which dominate at earlier stages of decay in the litter layer (Lindahl et al., 2007; Lu and Scheu, 2021). Investigating variations in stable isotope values with soil depth may provide deeper insight into opportunistic feeding and trophic niche differentiation of Oribatida species.

Similar to depth, earlier studies suggested that trophic niches of Oribatida vary little with forest type (Schneider et al., 2004). However, forest types vary in litter quality and microbial communities (Albers et al., 2004; Lu and Scheu, 2021), and different resource availability across forest types may induce changes in the use of basal resources and trophic positions of Oribatida species (Krause et al., 2019; Maraun et al., 2020). To date, only a few Oribatida species have been shown to respond flexibly by adjusting their trophic positions across land-use systems, but no agreement has been reached why some Oribatida species change their trophic niches with forest type while others do not (Gan et al., 2014; Krause et al., 2019; Maraun et al., 2020). The contrasting response calls for more detailed analyses including a wide range of Oribatida species with different traits across ecosystems. Guilds are groups of species that use similar class of resources and may differ in response to environmental conditions (Simberloff and Dayan, 1991). Including guilds of Oribatida may allow deeper insight into whether the variation of trophic niches in Oribatida species between forest types differ among guilds, e.g., is more pronounced in species at the bottom of the food web confronted with basal resources of different quality.

A better understanding of trophic niche variation in Oribatida species has important implications for forest management. In temperate and boreal regions, tree species richness is low, and managed forests are typically dominated by only one or few tree species (Knoke et al., 2008; Bauhus et al., 2010). The climax tree species in Central Europe is European beech (Leuschner et al., 2017), and litter decomposition in European beech forests is faster than e.g., in Norway spruce forests (Albers et al., 2004). Norway spruce, the economically most important timber species in Central Europe, is at increasing risk due to climate change and associated bark beetle outbreaks (Pettit et al., 2020). Although the admixture of Norway spruce to native European beech forests may reduce the risk of damage while maintaining economic gains, non-native Douglas fir is increasingly planted (Schmid et al., 2014). Recent studies indicated that the impact of Douglas fir on soil microbial communities is detrimental at nutrient-poor sites but not at nutrient-rich sites (Lu and Scheu, 2021), and Douglas fir monocultures change the composition of Oribatida communities (J.-Z. Lu, unpubl. data). Overall, the effects of forest type including non-native tree species and mixed forests on the functioning of the soil food webs are little studied (Schmid et al., 2014; Kriegel et al., 2021). The variation in trophic niches of Oribatida species across forest types may reflect changes in resource availability of different forest plantations, allowing deeper insight into linkages between tree species composition and belowground biota (Wardle et al., 2004).

Here, we quantified trophic niches of 40 Oribatida species in litter and soil of different forest types using bulk stable isotope analysis of ^13^C and ^15^N. Five forest types were investigated, including pure stands of European beech, Norway spruce, Douglas fir, and two conifer–beech mixtures. Forest types were replicated covering a range of water and soil nutrient conditions. We hypothesized that (1) stable isotope ratios of Oribatida species vary between litter and soil, and (2) variation in stable isotope ratios with soil depth is more pronounced in Oribatida species of low than in those of high trophic level. Further, we hypothesized that (3) particularly Oribatida species of low trophic level occupy different trophic niches across forest types, with the differences being most pronounced between European beech and coniferous forests.

## Methods

### Study sites

The study was conducted in 40 forest stands located in Northern Germany (8 sites x 5 forest types). The forest sites covered a wide range of soil nutrient and water conditions. Four more southern sites stock on Cambisol and Luvisol, with mean annual precipitation of 821–1029 mm. Four more northern sites are located on nutrient-poor out-washed sand with Podzol soil, with mean annual precipitation of 672–746 mm. Each site comprised three pure stands of European beech (*Fagus sylvatica* L.), Norway spruce (*Picea abies* [L.] Karst.), and Douglas fir (*Pseudotsuga menziesii* (Mirbel) Franco.) as well as two beech–conifer mixtures (European beech/Douglas fir and European beech/Norway spruce). Focal tree species in pure stands on average comprised more than 90% of total basal area, while in mixed stands, focal tree species accounted on average 34% for European beech and 58% for Douglas fir, and 56% for European beech and 37 % Norway spruce. Trees were on average more than 50 years old. More details on the sites are given in Ammer et al. (2020), Foltran et al. (2020), and Lu & Scheu (2021).

### Soil sampling

Animals were sampled by using a metal corer (ø 20 cm) between November 2017 and January 2018. Samples were taken between trees of the same (pure stands) or different species (mixed stands). One core was taken in each plot, and samples were separated into litter and 0–5 cm soil depth. Soil arthropods were extracted separately for litter and soil using high-gradient heat extraction (Kempson et al., 1963). Animals were collected in 50 % diethylene glycol and then transferred into 70 % ethanol. Species were identified using the key of Weigmann (2006). Separate soil samples (ø 5 cm) were taken in close vicinity, and were separated into litter and 0–5 cm soil for bulk stable isotope analysis (Lu and Scheu, 2021).

### Species selection

We collected species that occurred both in litter and soil from the same soil core, as our primary focus was to understand the variation in stable isotope values of Oribatida species with soil depth. From each soil core, two to three pairs of species (adults) were selected for stable isotope analysis. We aimed at covering a wide range of species, which were selected a priori based on a previous study at the same sites to cover representative trophic guilds (Lu et al. 2021, in prep.). Females were used in species with sexual dimorphism, i.e., *Acrogalumna longipluma* and *Adoristes ovatus*. In total, we analyzed 40 species including primary decomposers, secondary decomposers, as well as endophagous and predator/scavenger species (Table S1). Each species on average occurred in two forest plots, on average four replicates per species. This allowed estimating trophic niche variation across forests by controlling for species identity in mixed-effects models (see below). Guilds were balanced in each of the forest types (Table S2), allowing to test for the interaction between forest type and trophic guild.

### Bulk stable isotopes

We used bulk stable isotopes (^13^C/^12^C and ^15^N/^14^N ratios) of Oribatida to quantify their trophic niches. Stable isotope values of bulk litter and soil were measured, and used as baseline for comparing Oribatida species across forest types (Klarner et al., 2013; Potapov et al., 2019). Different numbers of individuals were used for stable isotope analysis depending on the body size of the species (Table S1), but for Oribatida species from different depths of the same soil core the same number of individuals was used. Animals, litter and soil were dried at 60°C for 48 h. Litter and soil were ground, and homogenized using a ball mill. After weighing samples into tin capsules, the natural abundance of stable isotope ratios of carbon (^13^C/^12^C) and nitrogen (^15^N/^14^N) of bulk litter and soil was determined by a coupled system of an elemental analyzer (NA 1110, CE-instruments, Rodano, Milano, Italy) and a mass spectrometer (Delta Plus, Finnigan MAT, Bremen, Germany). Isotopic signatures of animals were analyzed similarly by a coupled system of an elemental analyzer (Flash 2000, Thermo Fisher Scientific, Cambridge, UK) and a mass spectrometer (Delta V Advantage, Thermo Electron, Bremen, Germany). Animal samples with a dry weight <100 µg were analyzed using a modified set up adopted for small sample size (Langel and Dyckmans, 2014). Atmospheric nitrogen and Vienna PeeDee belemnite were used as primary standards. Acetanilide (C_8_H_9_NO, Merck, Darmstadt) was used as internal working standard. Natural variation in stable isotope ratios of carbon and nitrogen (δX) is expressed as δX (‰) = (R_sample_ - R_standard_) / R_standard_ Х 1000, with R being the ratio between the heavy and light isotopes (^13^C/^12^C or ^15^N/^14^N).

### Data analysis

We used linear mixed-effects models (LMMs) to analyze the variation in δ^13^C and δ^15^N values of Oribatida species. Fixed effects included depth (litter and 0–5 cm soil), forest type (European beech, Douglas fir, Norway spruce, mixture of European beech with Douglas fir, mixture of European beech with Norway spruce), trophic guild (primary decomposer, secondary decomposer, endophagous, predatory/scavenging), and site condition (nutrient-rich and nutrient-poor sites). We included several two-factor interactions (depth x forest type, depth x site condition, depth x trophic guild, and forest type x trophic guild) as fixed effects because our primary focus was (1) to evaluate depth effects and their dependencies on other factors, and (2) to evaluate forest type effects and their dependence on Oribatida guilds. The stable isotope values of litter and of 0–5 cm soil were included as covariates to control for differences in the baseline across forests (Melguizo-Ruiz et al., 2017). The cross-random effects included 40 plots and 40 species, accounting for the non-independence of samples from the same soil core, and for the taxonomic identity analyzed across forests.

To confirm the trophic guilds that were assigned a priori, we calibrated stable isotopes of Oribatida species using the stable isotope values of litter from the same plot (Scheu and Falca, 2000; Schneider et al., 2004; Klarner et al., 2013). In addition, we also modeled the variation in stable isotope values of bulk material as a function of forest type (European beech, Douglas fir, Norway spruce, European beech/Douglas fir, European beech/Norway spruce), depth (litter and 0–5 cm soil), site condition (nutrient-rich and nutrient-poor), and their interactions.

For Oribatida, we applied contrasts to inspect differences in their δ^13^C and δ^15^N values between litter and 0–5 cm soil depth. The contrast was designated as the difference of estimated marginal means between litter and soil (Piovia-Scott et al., 2019). We estimated the contrast for each forest type and each trophic guild. For litter and soil, we similarly applied contrasts to compare depth differences in δ^13^C and δ^15^N values of bulk litter and soil for each forest type. Furthermore, to quantify differences in δ^13^C and δ^15^N values of Oribatida species between forest types, we compared δ^13^C and δ^15^N values of Oribatida between forest types using European beech, the climax tree species in Central Europe, as reference (Leuschner et al., 2017; Lu and Scheu, 2021).

All analyses were done in R version 4.0.3 (https://www.r-project.org/). We used ‘lme4’ to fit LMMs (lmer) (Bates et al., 2015), and ‘emmeans’ to estimate marginal means. The package ‘lmerTest’ was used to derive P-values of LMMs with degrees of freedom estimated by Satterthwaite’s method (Kuznetsova et al., 2017). All LMMs met the assumptions of normality of residuals and homogeneity of variance.

## Results

The difference in δ^13^C and δ^15^N values between bulk litter and soil was highest in European beech (1.64 ± 0.23 ‰ and 3.85 ± 0.34 ‰ for δ^13^C and δ^15^N, respectively) and lowest in Douglas-fir forests (0.52 ± 0.23 ‰ and 2.57 ± 0.34 ‰ for δ^13^C and δ^15^N, respectively). Despite differences in depth-gradients of bulk stable isotope values in the studied forests (Table S3, Fig. S1), δ^13^C and δ^15^N values of Oribatida species did not differ significantly between litter and soil (Table 1, Fig. 1). The selected 40 species covered primary decomposers (n = 7), secondary decomposers (n = 17), endophagous (n = 9), and predatory/scavenging species (n = 7) (Table S1, Figs 2, S2). Trophic guilds explained the majority of the variability in δ^13^C and δ^15^N values of Oribatida species (in both cases ca. 56 % out of 64 % of total variation explained). However, δ^13^C and δ^15^N values of trophic guilds of Oribatida did not significantly vary with soil depth (Fig. 3).

**Table 1.** Linear mixed-effects models on δ^13^C and δ^15^N values of Oribatida species (Type III ANOVA). Fixed effects include Depth (litter and soil), Forest type (European beech, Douglas fir, Norway spruce and mixed forests of European beech and Douglas fir and European beech and Norway spruce), Guild (primary decomposer, secondary decomposer, endophagous, predatory), Site condition (nutrient-rich and nutrient-poor sites), and interactions of depth with other categorical factors. δ values of bulk litter and soil were included as covariates to control for the variation in stable isotope values of basal resources across forest ecosystems. Random effects included 40 plots and 40 species. Satterthwaite’s method was used to estimate denominator degrees of freedom (df). Significant P-values are in bold (P ≤ 0.05).

| Factor | $\delta^{13}\text{C}$ Oribatida | | | | - | $\delta^{15}\text{N}$ Oribatida | | | |
| --- | --- | --- | --- | --- | --- | --- | --- | --- | --- |
|  | df | SumSq | F | P |  | df | SumSq | F | P |
| Depth (D) | 1,94 | 0.004 | 0.022 | 0.883 |  | 1,93 | 2.393 | 2.121 | 0.149 |
| Forest type (F) | 4,35 | 4.258 | 6.097 | **0.001** |  | 4,27 | 6.460 | 1.432 | 0.251 |
| Guild (G) | 3,29 | 14.391 | 27.475 | **0.000** |  | 3,30 | 109.025 | 32.222 | **<0.001** |
| Site condition (S) | 1,37 | 0.850 | 4.867 | **0.034** |  | 1,25 | 0.316 | 0.280 | 0.601 |
| Soil Delta C | 1,25 | 0.072 | 0.411 | 0.527 |  | 1,22 | 1.052 | 0.932 | 0.345 |
| Litter Delta C | 1,27 | 0.101 | 0.578 | 0.454 |  | 1,25 | 14.595 | 12.940 | **0.001** |
| D x F | 4,93 | 0.490 | 0.702 | 0.592 |  | 4,92 | 5.855 | 1.298 | 0.277 |
| D x G | 3,93 | 0.075 | 0.143 | 0.934 |  | 3,93 | 3.267 | 0.965 | 0.413 |
| D x S | 1,93 | 0.210 | 1.201 | 0.276 |  | 1,92 | 2.518 | 2.232 | 0.139 |
| F x G | 12,111 | 3.690 | 1.761 | 0.063 |  | 12,127 | 16.743 | 1.237 | 0.265 |

**Figure 1.**
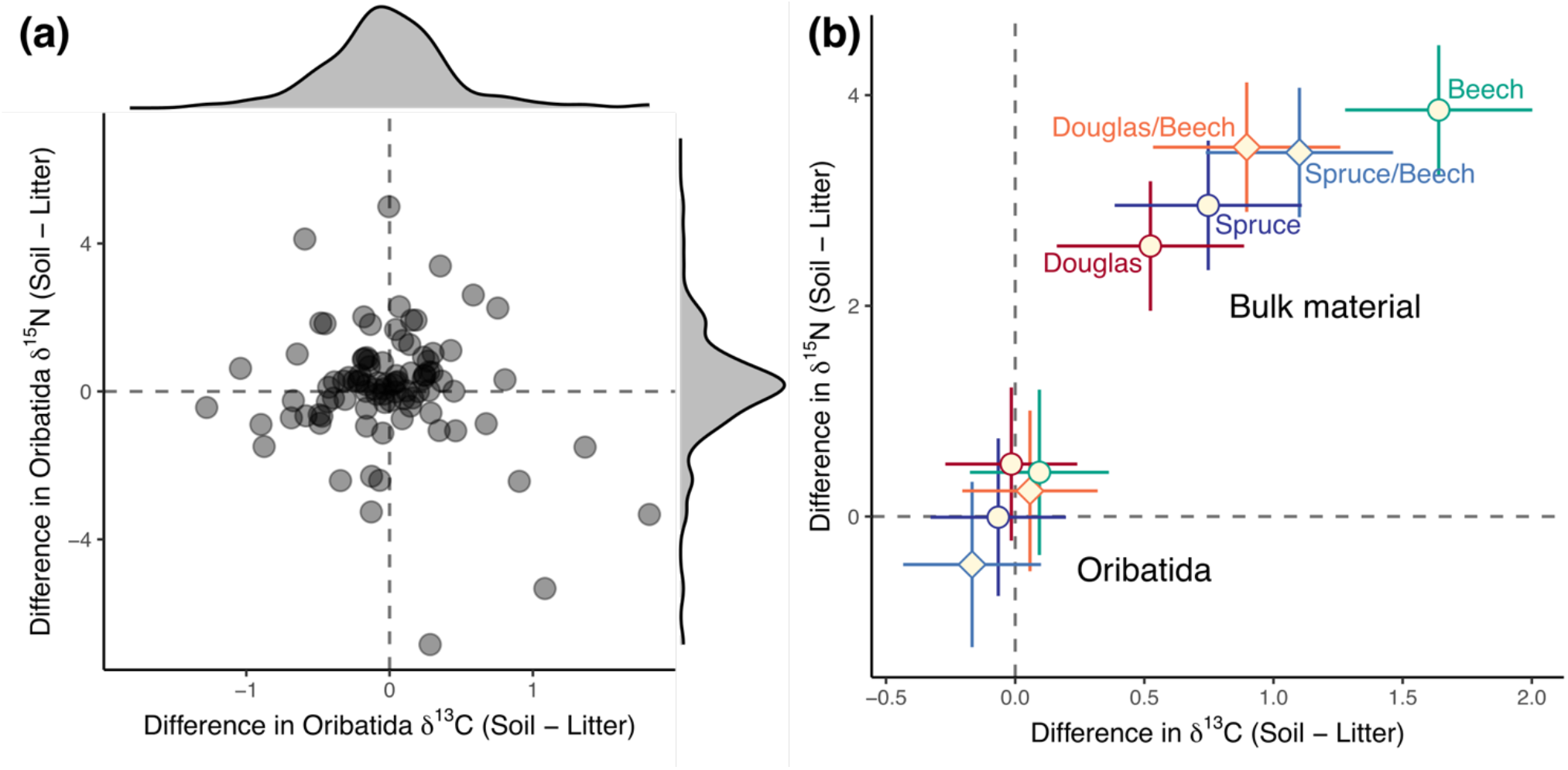
**(a)** Differences in δ^13^C and δ^15^N values of Oribatida species between soil and litter. **(b)** Differences in δ^13^C and δ^15^N values of Oribaitida and bulk soil between soil and litter in five forest types [European beech (Beech, green), Norway spruce (Spruce, blue), Douglas fir (Douglas, red), mixture of Norway spruce and European beech (Spruce/Beech, light-blue), mixture of Douglas fir and European beech (Douglas/Beech, light-red)]; means and 95% confidence intervals.

**Figure 2.**
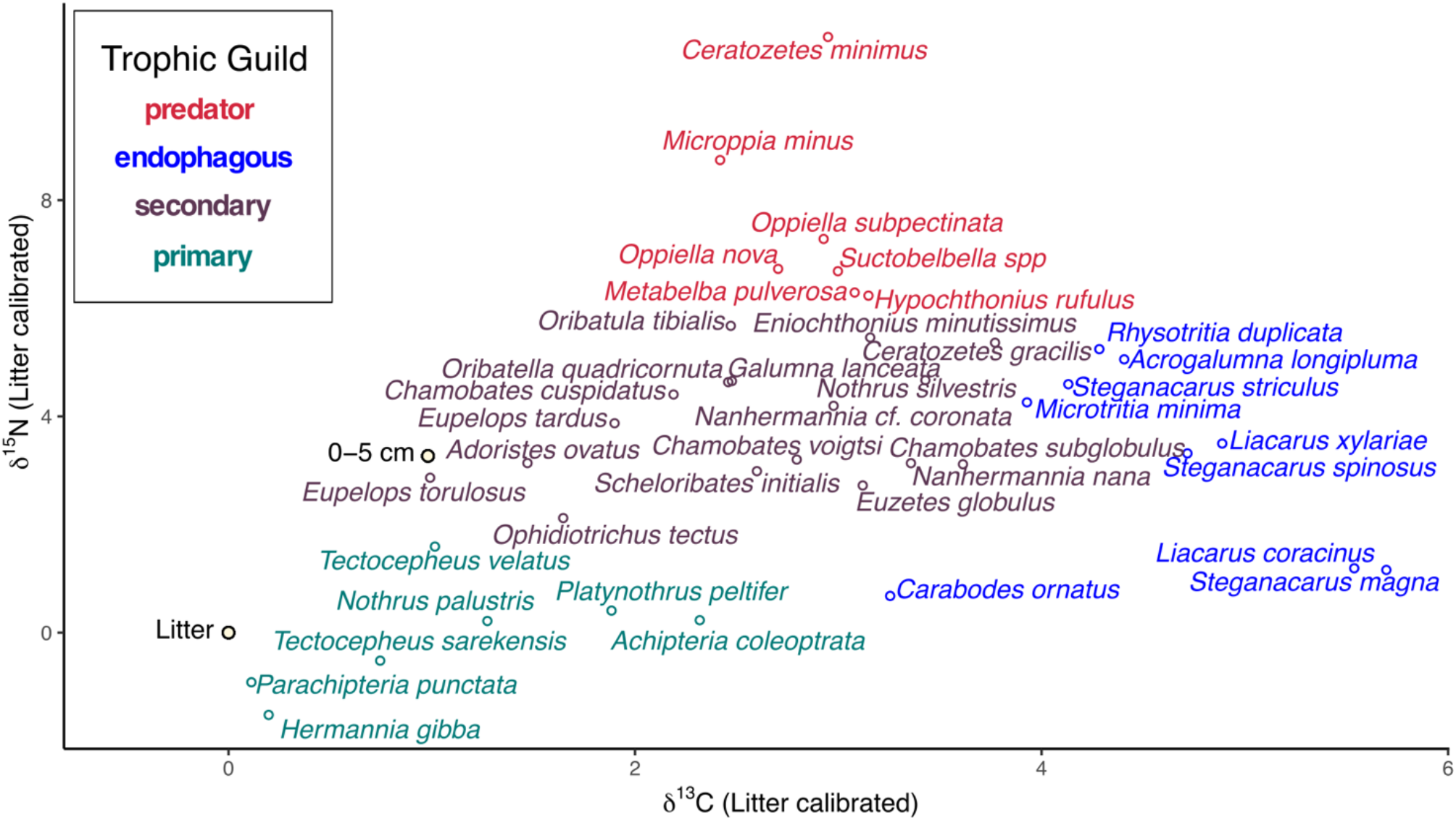
Trophic guilds of Oribatida species assigned based on litter calibrated δ^13^C and δ^15^N values. Means of Oribatid species and bulk material of litter and soil are shown; trophic guilds are color coded [primary decomposer (green), secondary decomposer (brown), endophagous (blue) and predatory (red)].

**Figure 3.**
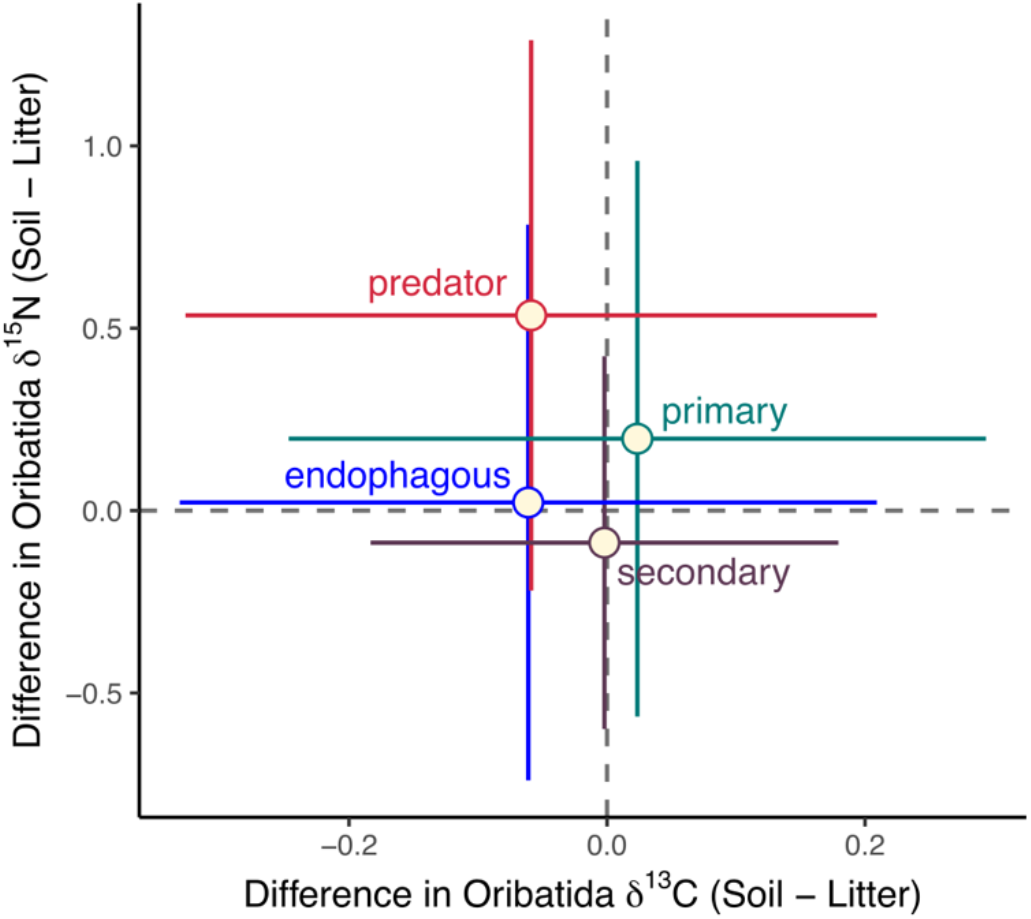
Differences in δ^13^C and δ^15^N values of trophic guilds of Oribatida [primary decomposer (green), secondary decomposer (brown), endophagous (blue) and predatory (red)]; means and 95% confidence intervals.

To account for variations in δ^13^C and δ^15^N values in basal resources across forest types, we included bulk stable isotope values of litter and 0–5 cm soil as covariates; stable isotope values of Oribatida species varied stronger with those of litter than with those of soil across forests (Table 1). Further, δ^13^C values of Oribatida species were higher in European beech than in Norway spruce (0.87) and Douglas-fir forests (0.58), and were intermediate in mixed forests (Figs 4, 5). The δ^13^C enrichment of Oribatida guilds in European beech forests was significant in secondary decomposer Oribatida, but not in primary decomposer and predatory/scavenging Oribaitda; in endophagous Oribaitda δ^13^C enrichment was higher in mixed forests than in European beech forests (Fig. S3). By contrast, as reflected by δ^15^N values, the trophic position of Oribatida species did not vary significantly between forest types (Fig. 4), and this was also true for each of the Oribatida guilds (Table 1).

**Figure 4.**
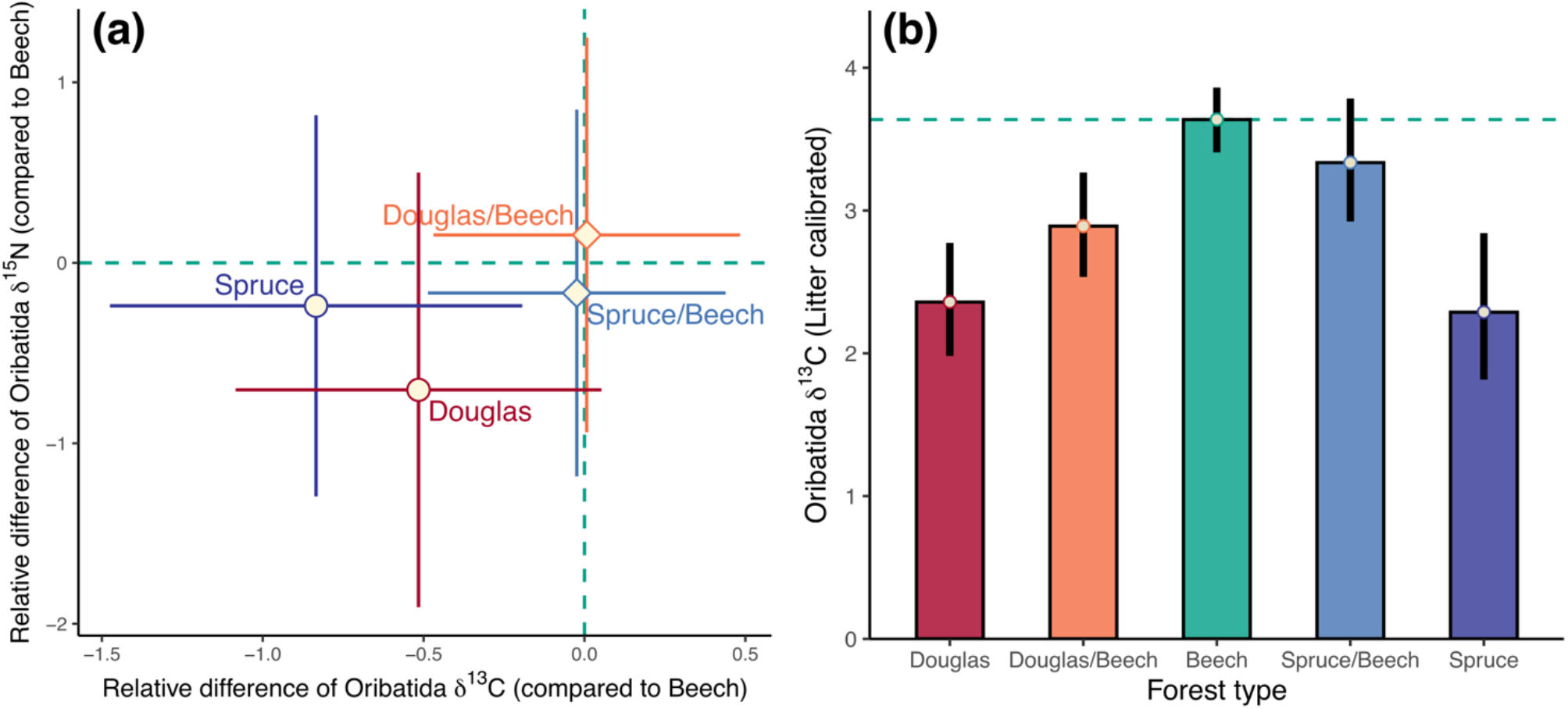
Trophic niche plasticity of Oribatida across forest types. **(a)** Difference in δ^13^C and δ^15^N values of Oribatida species in comparison to European beech forests (Beech, green); means and 95% confidence intervals. **(b)** Comparison of δ^13^C values of Oribatida species between forest types [European beech (Beech, green), Douglas fir (Douglas, red), Norway spruce (Spruce, blue), Douglas fir and European beech mixture (Spruce/Beech, light-red) and Norway spruce/European beech mixture (Spruce, light-blue)]; means and standard errors.

**Figure 5.**
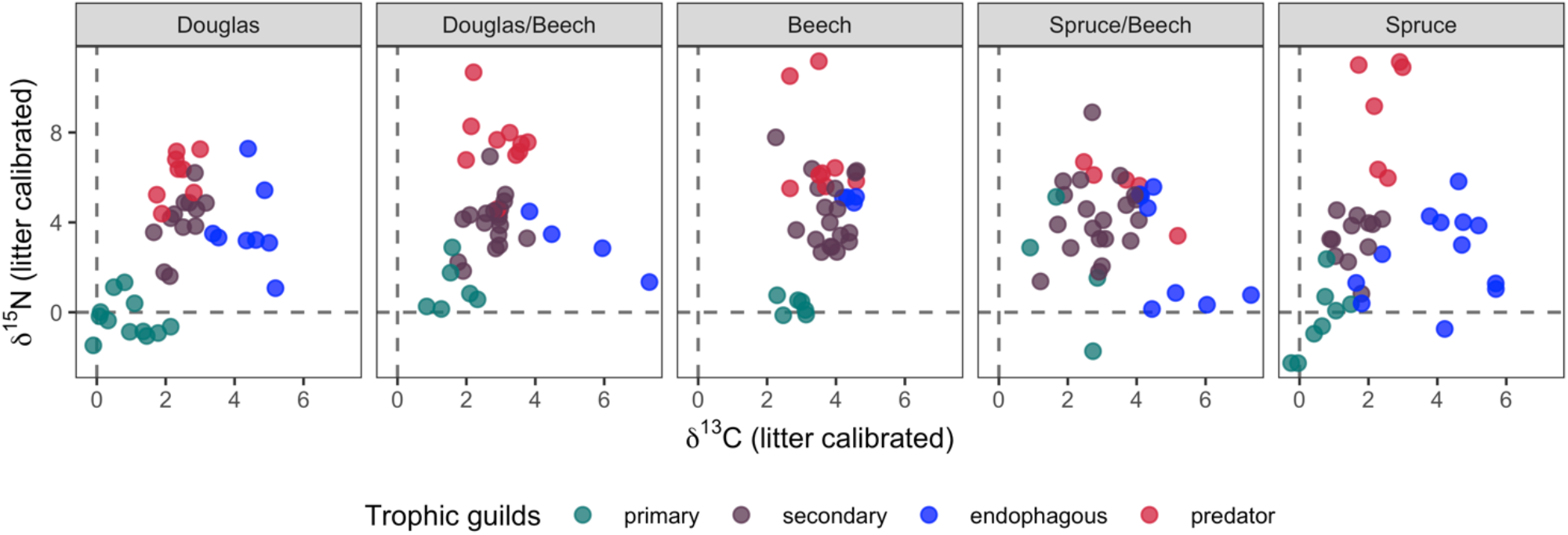
Litter calibrated δ^13^C and δ^15^N values of Oribatida guilds across forest types [Douglas fir (Douglas), mixture of Douglas fir/European beech (Douglas/Beech), European beech (Beech), mixture of Norway spruce/European beech (Spruce/Beech), Norway spruce (Spruce)]. Colors code for trophic guilds: primary decomposer (green), secondary decomposer (brown), endophagous (blue) and predatory species (red).

## Discussion

Trophic niche differentiation of soil microarthropods has advanced our understanding of species co-existence (Schneider et al., 2004), and low intraspecific variation may further strengthen species niche differentiation. To better understand the intraspecific variation in trophic niches of Oribatida, we investigated if trophic niches of Oribatida species vary with soil depths and forest types. Based on 40 Oribatida species occurring in litter and soil of temperate forests, we found that trophic niches of Oribatida species in litter and soil are highly consistent, and their trophic positions are similar across deciduous and coniferous forests. This implies that intraspecific variability in trophic niches of Oribatida is low, further facilitating the co-existence of soil Oribatida species (Hart et al., 2016).

### Variation with soil depth

Despite strong gradients in stable isotope values in bulk materials of the studied forest with soil depth, especially in European beech, trophic niches of Oribatida species were consistent irrespective of sampling depth. This rejects our first hypothesis and suggests that the actual environments of Oribatida species at measured scale do not inform us about where the majority of food resources have been acquired during their lifetime (Scheu and Falca, 2000). We sampled soil down to 5 cm depth, equivalent to ca. 50–200 times the body length of Oribatida, but Oribatida may be more mobile than commonly assumed (Åström and Bengtsson, 2011). Basal resources and trophic position of Oribatida species in litter and soil are highly consistent as indicated by δ^13^C and δ^15^N values of Oribatida species, despite that the decomposition rate and speed of internal nitrogen cycling differ between European beech and coniferous forests. Our results suggest that Oribatida may have access to their required resources irrespective of the soil depth they colonize. Further, stable isotope ratios in litter well explained the variation in stable isotope values of Oribatida species, suggesting that Oribatida species predominantly rely on fresh organic carbon in litter or on root-derived resources irrespective of the actual depth they inhabit (Pollierer et al., 2007; Okuzaki et al., 2009; Melguizo-Ruiz et al., 2017; Potapov et al., 2019; Pollierer and Scheu, 2021).

The lack of variation in trophic niches of Oribatida species with soil depth also applied to trophic guilds, contrasting our second hypothesis. In temperate forests, the relative availability of food resources, such as litter, bacteria, ectomycorrhizal, and saprotrophic fungi, changes rapidly with soil depth, with increasing proportions of bacteria and ectomycorrhizal fungi deeper in soil (Lindahl et al., 2007; Lu and Scheu, 2021). We investigated four Oribatida guilds typically occurring in temperate forests, spanning from primary and secondary decomposers, endophagous species to predators/scavengers (Schneider et al., 2004; Maraun et al., 2011). As trophic guilds of Oribatida differ in food resources (Maraun et al., 2011), primary decomposer and predatory Oribatida, for example, differ most in trophic position, they may respond differently to soil depth. However, our results strongly suggest that Oribatida guilds occupy similar niches irrespective of soil depth, further supporting that despite resource quantity and quality change with soil depth (Lu and Scheu, 2021), the trophic niches of Oribatida remain unaffected (Scheu and Falca, 2000). Notably, this applies to a wide range of species (n = 40) including major trophic guilds of Oribatida (n = 4), colonizing pure and mixed forests of deciduous and coniferous trees (n = 5) arguing for the generality of these results.

The lack of variation in stable isotope values with soil depth in Oribatida species may point towards the high vertical mobility of these species; however, it also points to the presence of these niches irrespective of soil depth in the forest floor. In any case, the high consistency of trophic niches in Oribatida species underlines the importance of niche differentiation for the co-existence of Oribatida species in soil (Schneider et al., 2004). Trophic niche consistency in Oribatida species also indicates that intraspecific competition may be stronger than interspecific competition, thereby facilitating niche differentiation of Oribatida species in soil (Hart et al., 2016). Although the contribution of bacteria and plants as basal resources to Oribatida species varies depending on species, saprotrophic fungi are the major food resource of primary and secondary decomposer Oribatida species as indicated by compound-specific stable isotope analysis of amino acids (Pollierer and Scheu, 2021). In contrast to different life forms in Collembola (Potapov and Tiunov, 2016), the vertical distribution of Oribatida species has not yet been found to be related to morphological and life-history traits (Pande and Berthet, 1975; Potapov et al., 2016). The mechanisms responsible for maintaining species diversity, therefore, are likely to differ between Collembola and Oribatida. Our findings also challenge the view that Oribatida species are opportunistic feeders (Maraun and Scheu, 2000), but rather Oribatida species occupy a distinct niche in the field irrespective of the depth and forest type they inhabit.

### Variation with forest type

The detrital shift, i.e., the enrichment in δ^13^C values relative to litter (Potapov et al., 2019), has been widely documented in terrestrial ecosystems (Pollierer et al., 2009; Potapov et al., 2019; Susanti et al., 2021), suggesting that it is a universal phenomenon in decomposer food webs. We found that the detrital shift is stronger in European beech than in coniferous forests. The contrasting detrital shift between European beech and coniferous forests suggests that the basal resources of Oribatida species differ between forest types, supporting our third hypothesis. Several non-mutually exclusive mechanisms may explain the contrasting detrital shift between deciduous and coniferous forests. The litter quality of European beech may be particularly poor and rich in lignin (depleted in ^13^C), and saprotrophic microorganisms incorporate more palatable litter compounds that are enriched in ^13^C compared to lignin (Pollierer et al., 2009). δ^13^C values of leaf litter of European beech (−29.58 ± 0.33 ‰) are indeed about 1.27–1.41 ‰ lower than those in needle litter of coniferous trees (−28.31 ± 0.37 ‰ and -28.17 ± 0.44 ‰ for Douglas fir and Norway spruce, respectively; mean ± sd); further, the C/N ratio in the leaf litter of European beech (50.73 ± 6.26) is higher than that in needle litter of coniferous trees (39.26 ± 1.85 and 42.3 ± 5.21 for Norway spruce and Douglas fir, respectively; J.-Z. Lu, unpubl. data), which is in agreement to earlier suggestions that spruce needles are not more recalcitrant than beech leaves and may even decompose faster (Albers et al., 2004; Berger and Berger, 2012). Notably, however, as indicated by microbial biomass and microbial basal respiration, microbial activity in European beech forests exceed that in coniferous forests, which may be due to more efficient decomposer communities and/or favorable abiotic conditions (Albers et al., 2004; Lu and Scheu, 2021). High microbial activity in European beech forests drives faster carbon turnover, and increases the incorporation of microbial processed carbon into soil food webs (Potapov et al., 2019), presumably contributing to the pronounced detrital shift in European beech forests. Supporting the importance of microbial activity, differences in the detrital shift between European beech and coniferous forests were strongest in secondary decomposer Oribatida, known as predominant fungal feeders (Schneider et al., 2004). Our findings on the detrital shift likely also apply to other soil invertebrates feeding on microorganisms in soil food webs (Scheu and Falca, 2000; Pollierer et al., 2009; Klarner et al., 2013).

Although basal resources differ between European beech and coniferous forests, Oribatida species kept their trophic position and this was true across Oribatida guilds and forest types, contrasting our third hypothesis. The paradox of opportunistic feeding at laboratory conditions while constant in trophic positions in the field call for a better understanding of the foraging ecology of Oribatida species. The importance of plant rhizosphere as microhabitats has been supported by a recent labeling experiment for Collembola (Li et al., 2021). Further, the high consistency of the trophic positions of Oribatida species supports that soil food webs are resistant to changes in forest types (Pollierer et al., 2021), agreeing with recent studies on variations in trophic niches in other mesofauna groups, such as Mesostigmata mites and Collembola between land-use systems (Klarner et al., 2017; Susanti et al., 2021). In tropical ecosystems ranging from rainforest to monoculture plantations, trophic plasticity of Oribatida has been suggested to depend on species (Krause et al., 2019), which we could not confirm for Oribatida species in temperate forest ecosystems. Adding to these findings, our results further suggest that, at least in temperate forests, trophic positions of Oribatida species does not vary among forest types, and the consistency also applies to Oribatida guilds.

High trophic level Oribatida species (i.e., predators/scavengers) on average were enriched in δ^15^N by 6.7–11.0 ‰, consistent with the enrichment of other predators in temperate forests including Mesostigmata mites, Chilopoda, and Araneida (Pollierer et al., 2009; Potapov et al., 2019). Incorporation of old organic carbon in detritivores may inflate their high trophic positions. However, the consistency in trophic positions of Oribatida species irrespective of soil depth indicates that tissue carbon in the diet of Oribatida is unlikely to be derived from old organic matter, rather it likely originates predominantly from litter and/or root-derived resources which are little decomposed (Klarner et al., 2013). If the detrital shift in Oribatida species would be due to the consumption of carbon derived from old organic matter, the shift should be stronger in soil than in litter, and Oribatida species should also be enriched in δ^13^C. Contrasting this assumption, δ^13^C and δ^15^N values of Oribatida species correlated more closely with those of litter than with those of soil. Therefore, we suggest that high trophic level Oribatida species live as predators/scavengers rather than incorporating nitrogen originating from old organic matter, contrasting other mesofauna with high δ^15^N values such as euedaphic Collembola (Potapov et al., 2019). However, some Oribatida species such as *Oppiella nova* has been suggested to feed on ectomycorrhizal fungi, known to be enriched in ^15^N (Remén et al., 2008; Potapov and Tiunov, 2016). Notably, *Oppiella nova* was the most abundant soil mesofauna species at our study sites, and the resources used by this species, therefore, need further attention including experiments manipulating the input of root-derived resources (Bluhm et al., 2021).

The high consistency in trophic positions of Oribatida guilds across soil depth and forest types support the incorporation of trophic guilds in soil food webs. Early studies analyzing natural variations in stable isotope ratios indicated Oribatida species to live as primary and secondary decomposers (Scheu and Falca, 2000), but later Oribatida species have been shown to cover virtually the full range of the isotope space of soil food webs (Schneider et al., 2004; Maraun et al., 2011). Despite that high δ^13^C in endophagous Oribatida may inflate their δ^13^C niche due to incorporation of calcium carbonate (Maraun et al., 2011), the isotopic ranges of Oribatida covered 100 % (δ7.56) of the δ^13^C and 67 % (δ13.43) of the δ^15^N of the isotopic range of the whole soil food webs (Potapov et al., 2019). Ascribing Oribatida species into guilds reduces the complexity of soil food webs (Nielsen, 2019) and has the potential to better represent their roles in soil food webs. Grouping Oribatida species into guilds reduce the complexity of soil food webs (Nielsen, 2019). Results of this study also call to for studying the depth variation of trophic niches in other soil animal groups such as Collembola and Gamasina to move toward more spatially explicit trophic interactions in soil food webs.

### Management implications

In face of climate change, Norway spruce is increasingly threatened in the lowland of Central Europe. Mixed forests and alternative tree species such as non-native Douglas fir may provide better options for the provisioning of long-term ecosystem services than monoculture plantations of Norway spruce. By studying the variation in trophic niches in an abundant group of soil microarthropods, we found that trophic positions of Oribatida are consistent across European beech and coniferous forests, suggesting that the trophic structure of Oribatida species is highly resistant against the plantation of different tree species including non-native species such as Douglas fir. Although basal resources of Oribatida vary between coniferous and deciduous forests, the effects of Douglas fir on basal resources of Oribatida species did not differ from that of Norway spruce. Further, the basal resources in mixed forests were similar to those in European beech forests, supporting the potential of mixed forests to mitigate detrimental effects of coniferous trees on ecosystem properties and functioning. Although to gain a holistic understanding of forest type effects on the functioning of the decomposer system more information is needed on the energy flux through other taxa of soil food webs, we conclude that trophic niches of Oribatida species and guilds are highly consistent in European beech and Douglas fir mixed forests.

## Authorship

JZL, SS and MM designed the study. JZL, PC collected the data. JZL analyzed the data and wrote the first draft of the manuscript. All authors contributed significantly to interpretations and revisions.

## Acknowledgements

We are indebted to S. Maurer, S. Böning-Klein, T. Volovei, C. Bluhm, S.-X. Zhou, R. Langel, and J. Dyckmans for assistant. We thank A. Potapov, I. Schaefer, H. Riebl, A. Krause, S. Meyer, and S. Bluhm for helpful discussions. Special thanks to S. Müller for coordinating the project (Enrico; RTG 2300). This work was supported by the German Research Foundation (Grant ID: 316045089).

