## supplemental figures and tables for "High consistency of trophic niches in soil microarthropod species (Oribatida, Acari) across soil depth and forest type"

### Supplementary Tables

**Table S1**

Species list Oribatida (n = 40). Trophic guilds were assigned according to litter calibrated $\delta^{13}C$ and $\delta^{15}N$ values: primary decomposer, secondary decomposer, endophagous Oribatida and scavenger/predator. Total number of animals for each species used for stable isotopes and their ranges (min - max) are given.

| **Oribatid taxa** | **Family** | **Total number (range)** | **$\delta^{13}C$** | **$\delta^{15}N$** | **Trophic guilds** |
|---|---|---|---|---|---|
| *Ceratozetes minimus* Sellnick, 1928 | Ceratozetidae | 10 (10-10) | 2.95 ± 0.06 | 11.02 ± 0.17 | predator |
| *Hypochthonius rufulus* C. L. Koch, 1835 | Hypochthoniidae | 4 (2-7) | 3.15 ± 0.77 | 6.23 ± 0.96 | predator |
| *Metabelba pulverosa* Strenzke, 1953 | Damaeidae | 3 (3-3) | 3.08 ± 0.25 | 6.29 ± 2.40 | predator |
| *Microppia minus* (Paoli, 1908) | Oppiidae | 19 (7-25) | 2.42 ± 0.28 | 8.74 ± 2.42 | predator |
| *Oppiella nova* (Oudemans, 1902) | Oppiidae | 14 (8-17) | 2.70 ± 1.84 | 6.73 ± 2.79 | predator |
| *Oppiella subpectinata* (Oudemans, 1900) | Oppiidae | 9 (3-16) | 2.93 ± 0.93 | 7.28 ± 1.96 | predator |
| *Suctobelbella spp* Jacot, 1937 | Suctobelbidae | 22 (18-26) | 3.00 ± 0.74 | 6.69 ± 0.72 | predator |
| *Acrogalumna longipluma* (Berlese, 1904) | Galumnidae | 4 (3-5) | 4.41 ± 0.18 | 5.06 ± 0.12 | endophagous |
| *Carabodes ornatus* Storkan, 1925 | Carabodidae | 2 (1-3) | 3.26 ± 1.79 | 0.68 ± 0.52 | endophagous |
| *Liacarus coracinus* (C. L. Koch, 1841) | Liacaridae | 1 (1-2) | 5.54 ± 1.92 | 1.19 ± 1.37 | endophagous |
| *Liacarus xylariae* (Schrandk, 1803) | Liacaridae | 2 (1-2) | 4.89 ± 0.26 | 3.50 ± 1.98 | endophagous |
| *Microtritia minima* (Berlese, 1904) | Euphthiracaridae | 12 (10-15) | 3.93 ± 0.56 | 4.26 ± 1.05 | endophagous |
| *Rhysotritia duplicata* (Grandjean, 1953) | Euphthiracaridae | 8 (8-9) | 4.28 ± 0.46 | 5.24 ± 1.49 | endophagous |
| *Steganacarus magnus* (Nicolet, 1855) | Phthiracaridae | 1 (1-1) | 5.7 ± 0 | 1.16 ± 0.18 | endophagous |
| *Steganacarus spinosus* (Sellnick, 1920) | Phthiracaridae | 4 (1-8) | 4.72 ± 0.36 | 3.32 ± 0.38 | endophagous |
| *Steganacarus striculus* (C. L. Koch, 1835) | Phthiracaridae | 4 (3-5) | 4.13 ± 0.26 | 4.60 ± 0.82 | endophagous |
| *Adoristes ovatus* (C. L. Koch, 1839) | Liacaridae | 1 (1-1) | 1.47 ± 0.58 | 3.14 ± 1.63 | secondary |
| *Ceratozetes gracilis* (Michael, 1884) | Ceratozetidae | 8 (6-10) | 3.77 ± 0.93 | 5.36 ± 1.03 | secondary |
| *Chamobates cuspidatus* (Michael, 1884) | Chamobatidae | 4 (2-7) | 2.19 ± 0.70 | 4.41 ± 0.43 | secondary |
| *Chamobates subglobulus* (Oudemans, 1900) | Chamobatidae | 1 (1-1) | 3.36 ± 0.57 | 3.13 ± 0.22 | secondary |
| *Chamobates voigtsi* (Oudemans, 1902) | Chamobatidae | 6 (2-9) | 2.80 ± 0.64 | 3.20 ± 1.49 | secondary |
| *Eniochthonius minutissimus* (Berlese, 1903) | Eniochthoniidae | 8 (2-16) | 3.16 ± 1.15 | 5.46 ± 0.88 | secondary |
| *Eupelops tardus* (C. L. Koch, 1835) | Phenopelopidae | 1 (1-1) | 1.90 ± 0.35 | 3.87 ± 0.44 | secondary |
| *Eupelops torulosus* (C. L. Koch, 1839) | Phenopelopidae | 1 (1-1) | 0.99 ± 0.06 | 2.87 ± 0.52 | secondary |
| *Euzetes globulus* (Nicolet, 1855) | Euzetidae | 1 (1-1) | 3.12 ± 1.12 | 2.72 ± 0.26 | secondary |
| *Galumna lanceata* (Oudemans, 1900) | Galumnidae | 2 (1-2) | 2.48 ± 0.32 | 4.65 ± 0.80 | secondary |
| *Nanhermannia cf. coronata* Berlese, 1913 | Nanhermanniidae | 8 (2-14) | 2.98 ± 0.91 | 4.20 ± 0.32 | secondary |
| *Nanhermannia nana* (Nicolet, 1855) | Nanhermanniidae | 3 (3-3) | 3.61 ± 0.76 | 3.12 ± 0.96 | secondary |
| *Nothrus silvestris* Nicolet, 1855 | Nothridae | 3 (1-5) | 3.43 ± 0.67 | 4.67 ± 1.56 | secondary |
| *Ophidiotrichus tectus* (Michael, 1884) | Oribatellidae | 6 (4-8) | 1.65 ± 0.62 | 2.12 ± 1.05 | secondary |
| *Oribatella quadricornuta* Michael, 1880 | Oribatellidae | 2 (2-3) | 2.46 ± 0.40 | 4.64 ± 1.07 | secondary |
| *Oribatula tibialis* (Nicolet, 1855) | Oribatulidae | 3 (1-5) | 2.47 ± 0.35 | 5.68 ± 1.99 | secondary |
| *Scheloribates initialis* (Berlese, 1908) | Scheloribatidae | 4 (3-6) | 2.60 ± 0.60 | 2.99 ± 0.78 | secondary |
| *Achipteria coleoptrata* (Linne, 1758) | Achipteriidae | 6 (1-10) | 2.32 ± 0.93 | 0.23 ± 1.19 | primary |
| *Hermannia gibba* (C. L. Koch, 1839) | Hermanniidae | 2 (2-2) | 0.20 ± 0.41 | -1.52 ± 0.86 | primary |
| *Nothrus palustris* C. L. Koch, 1839 | Nothridae | 1 (1-1) | 1.27 ± 0.31 | 0.22 ± 0.20 | primary |
| *Parachipteria punctata* (Nicolet, 1855) | Achipteriidae | 2 (2-2) | 0.11 ± 0.30 | -0.92 ± 0.78 | primary |
| *Platynothrus peltifer* (C. L. Koch, 1839) | Camissidae | 2 (1-2) | 1.88 ± 0.66 | 0.41 ± 0.38 | primary |
| *Tectocepheus sarekensis* Traegardh, 1910 | Tectocepheidae | 15 (15-15) | 0.75 ± 0.75 | -0.52 ± 0.51 | primary |
| *Tectocepheus velatus* (Michael, 1880) | Tectocepheidae | 11 (5-19) | 1.02 ± 0.53 | 1.59 ± 1.85 | primary |

**Table S2**

Cross tabulation summarizing the study design. Frequency of trophic guilds (primary decomposer, secondary decomposer, endophagous and predator Oribatida) in each of the forest type [Douglas fir (Douglas), Douglas fir/European beech (Douglas/Beech), European beech (Beech), Norway spruce/European beech (Spruce/Beech), Norway spruce (Spruce)].

| | Douglas | Douglas/Beech | Beech | Spruce/Beech | Spruce |
|---|---|---|---|---|---|
| primary | 12 | 6 | 6 | 4 | 8 |
| secondary | 12 | 18 | 18 | 20 | 12 |
| endophagous | 8 | 4 | 4 | 8 | 12 |
| predator | 8 | 10 | 8 | 6 | 6 |

**Table S3**

Linear mixed-effects models on $\delta^{13}C$ and $\delta^{15}N$ values of bulk soil and litter (Type III ANOVA). Fixed effects include Depth (litter and soil), Forest type (European beech, Douglas fir, Norway spruce and mixed forests of European beech and Douglas fir and European beech and Norway spruce), Site condition (nutrient-rich and nutrient-poor sites), and their interactions. Random effects included 40 plots. Satterthwaite's method was used to estimate denominator degrees of freedom (df). Significant P-values are in bold ($P \leq 0.05$).

| | $\delta^{13}$ C Bulk Material | | | | | $\delta^{15}$ N Bulk Material | | | |
|---|---|---|---|---|---|---|---|---|---|
| Factor | df | SumSq | F | P | - | df | SumSq | F | P |
| Depth (D) | *1,30* | 19.261 | 152.873 | **<0.001** | | 1,30 | 213.667 | 589.591 | **<0.001** |
| Forest type (F) | *4,30* | 3.146 | 6.242 | **0.001** | | 4,30 | 9.061 | 6.251 | **0.001** |
| Site condition (S) | *1,30* | 0.000 | 0.000 | 0.997 | | 1,30 | 24.091 | 66.476 | **<0.001** |
| D x F | *4,30* | 2.877 | 5.708 | **0.002** | | 4,30 | 4.123 | 2.844 | **0.041** |
| D x S | *1,30* | 2.879 | 22.851 | **<0.001** | | 1,30 | 3.083 | 8.507 | **0.007** |
| F x S | *4,30* | 0.390 | 0.774 | 0.551 | | 4,30 | 1.734 | 1.196 | 0.333 |
| D x F x S | *4,30* | 0.619 | 1.227 | 0.32 | | 4,30 | 1.799 | 1.241 | 0.315 |

### Supplementary Figures

**Figure S1**

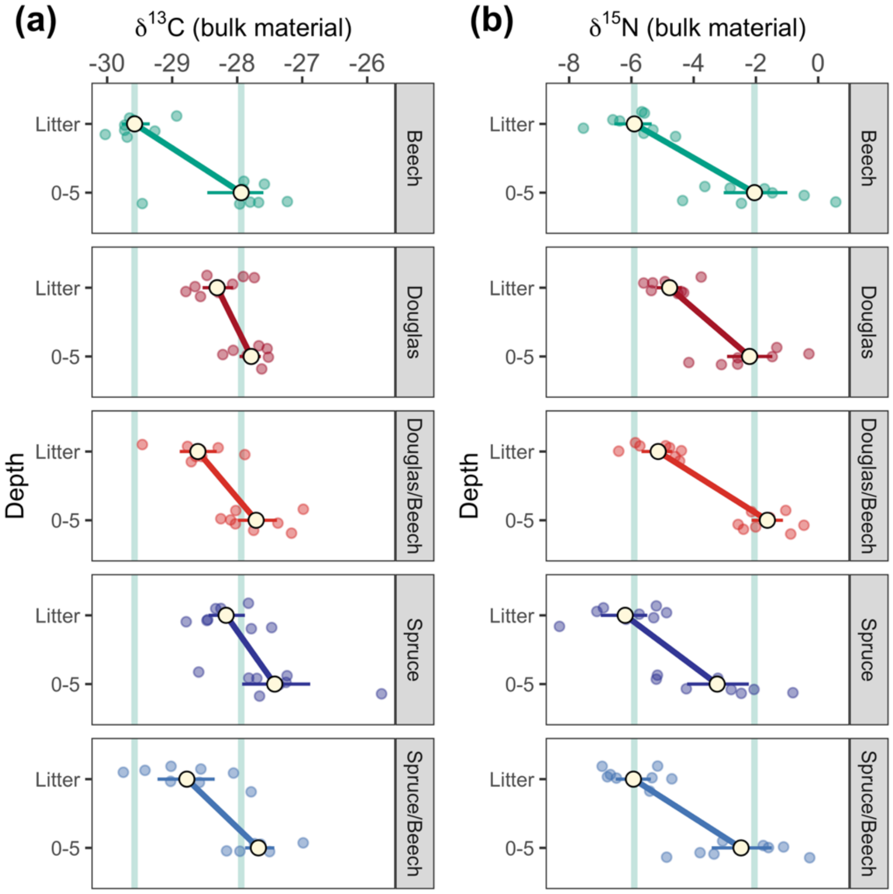


**Figure S1**. Stable isotope values of $\delta^{13}C$ (a) and $\delta^{15}N$ (b) of bulk litter and 0–5 cm soil in European beech (Beech), Douglas fir (Douglas), Douglas fir/European beech (Douglas/Beech), Norway spruce (Spruce), Norway spruce/European beech (Spruce/Beech) forests. Horizontal bars are bootstrap estimated standard errors (n = 8). The green vertical bars represent respective values in beech forests in litter and 0–5 cm soil.

**Figure S2**

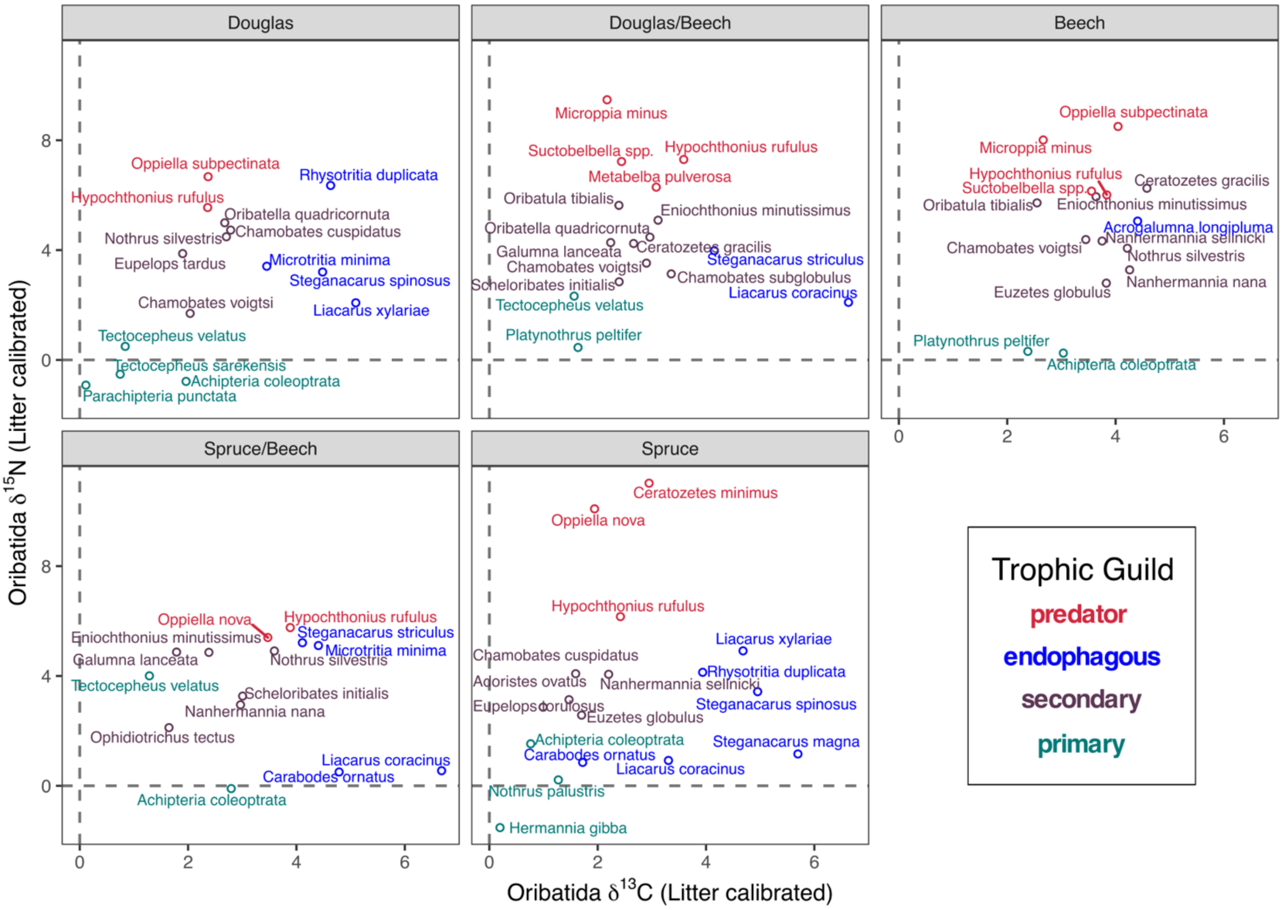


**Figure S2**. Average of litter calibrated δ¹³C and δ¹⁵N values of Oribatida species in each of the forest type [Douglas fir (Douglas), mixture of Douglas fir/European beech (Douglas/Beech), European beech (Beech), mixture of Norway spruce/European beech (Spruce/Beech), Norway spruce (Spruce)]. Colors code for trophic guilds: primary decomposer (green), secondary decomposer (brown), endophagous (blue) and predatory species (red).

**Figure S3**

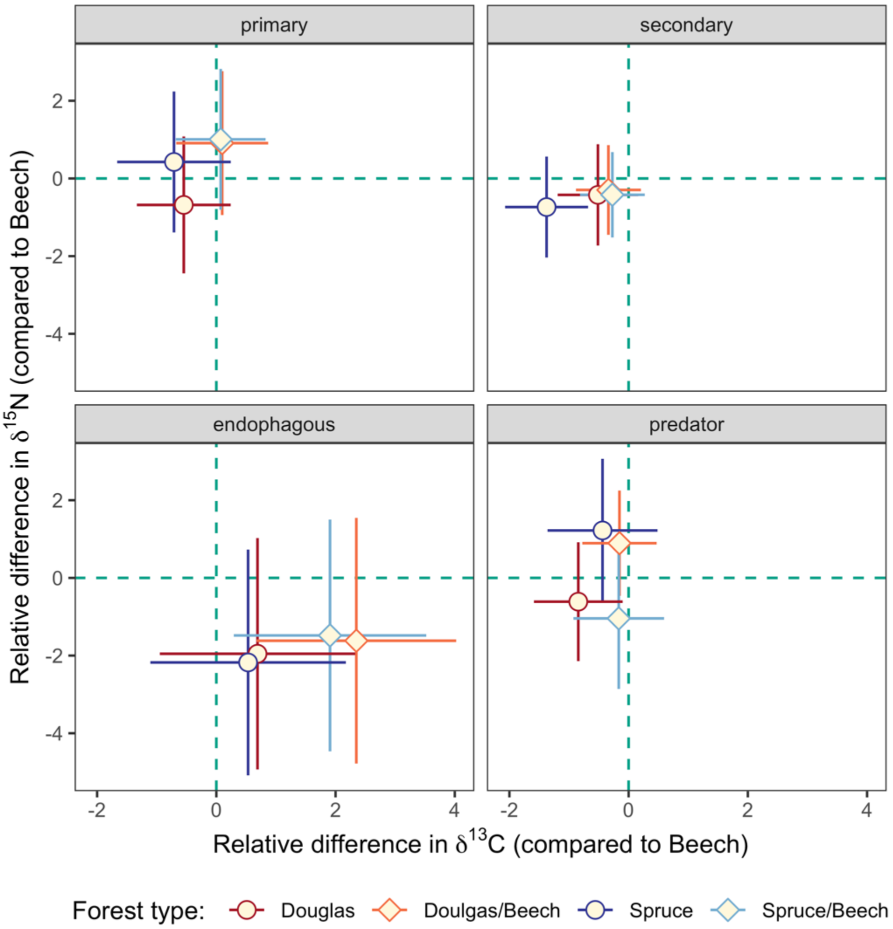


**Figure S3**. Difference in $\delta^{13}C$ and $\delta^{15}N$ values of Oribatida guilds (primary decomposer, secondary decomposer, endophagous and predatory) in comparison to European beech forests (Beech, dash line in green); Forest types include Douglas fir (Douglas, red), Norway spruce (Spruce, blue), Douglas fir and European beech mixture (Spruce/Beech, light-red) and Norway spruce/European beech mixture (Spruce, light-blue); means and 95% confidence intervals.
